# Spatial Transcriptomics Reveals *CXCL12*⁺ Fibroblasts as Central Immune Organizers through CXCR4 Signaling in Abdominal Aortic Aneurysm

**DOI:** 10.1101/2025.11.24.690328

**Authors:** Dina Levy-Lambert, Joel L. Ramirez, Saba Shaikh, Cesar De Jeronimo Diaz, April G. Huang, Alexis Combes, Gabriela K. Fragiadakis, Trevor P. Fidler, Adam Z. Oskowitz

## Abstract

**BACKGROUND:** Abdominal aortic aneurysm (AAA) is characterized by sterile inflammation, immune cell infiltration, and stromal remodeling that progressively weaken the aortic wall, leading to life-threatening aortic rupture. The molecular mechanisms and spatial organization of immune–stromal interactions in human tissue are poorly understood, limiting the potential to develop effective pharmacological therapy for AAA.

**METHODS:** In this observational cross-sectional study, formalin-fixed, paraffin-embedded tissues from 11 AAA patients and 12 controls were analyzed by Xenium spatial transcriptomics. Cellular states and localization within tissue architecture were mapped to identify cellular neighborhoods and infer cell–cell communication.

**RESULTS:** We generated a high-resolution spatial transcriptomics atlas of 581,664 cells in 26 clusters. AAAs showed a significant loss of contractile smooth muscle cells, expansion of pro-angiogenic endothelial subsets, and broad infiltration of immune cells. These inflammatory changes were accompanied by expansion of activated, universal, and *CXCL12⁺* adventitial fibroblasts. Spatial transcriptomic analysis revealed fibroblast–immune colocalization and adventitial tertiary lymphoid organs. Inferred signaling pathway analysis identified increased interactions between *CXCL12*⁺ fibroblasts and *CXCR4*⁺ T and B cells in the adventitia of AAAs. Fibroblasts that expressed *CXCL12* had significantly more immune cell neighbors than fibroblasts that did not, suggesting that they serve as stromal hubs for adaptive immune clustering. Genome-wide association analysis linked AAA heritability to fibroblasts, modulated smooth muscle cells, and foamy macrophages.

**CONCLUSION:** Our novel high-resolution spatial transcriptomic atlas of human AAAs revealed coordinated pathogenic reprogramming of stromal and immune cells, defined by smooth muscle cell depletion, fibroblast activation, endothelial remodeling, and disproportionate expansion of immune cells. Through *CXCR4* signaling, *CXCL12*⁺ fibroblasts serve as central organizers of immune niches, suggesting stromal–immune crosstalk as a therapeutic target in AAA.

**CLINICAL PERSPECTIVES:** What Is New?

- We generated the first subcellular-resolution spatial transcriptomic atlas of human abdominal aortic aneurysm (AAA), with >580,000 cells identified from aortic tissue sections
- We identified CXCL12^+^ fibroblasts as central stromal hubs that organize adaptive immune niches through CXCR4-mediated crosstalk with B and T cells
- We discovered that stromal populations carry the strongest genetic enrichment for AAA risk, notably fibroblast and modulated smooth muscle cell populations

What Are The Clinical Implications?

- These findings position stromal-immune interactions, particularly the CXCL12-CXCR4 axis, as a potential therapeutic target to slow AAA progression
- The spatial atlas provides a framework for mechanistic studies and drug-discovery efforts, guiding future interventions aimed at modifying the microenvironment that destabilizes the aneurysmal aortic wall

## INTRODUCTION

Abdominal aortic aneurysm (AAA) affects 4–9% of men and 1% of women over 65 years old and carries a risk of fatal rupture.^1^ Extracellular matrix (ECM) degradation, smooth muscle cell (SMC) apoptosis, fibroblast activation, neovascularization, and immune cell infiltration contribute to AAA pathology by weakening the vessel wall.^2–6^ The only definitive treatment is surgical repair, which entails perioperative mortality of 1–4%.^7,8^ Deeper insights into AAA pathogenesis are needed to develop pharmacologic treatment options.^9–12^

Transcriptomic assays have implicated specific cell types in AAA pathogenesis. A single-cell RNA sequencing (scRNA-seq) study of human AAAs reported clonal expansion of T and B lymphocytes, heterogeneity of myeloid subsets, and modulation of SMC phenotypes, including dedifferentiation, loss of contractile markers, and increased expression of genes involved in ECM remodeling and inflammation.^17^ Cell–cell communication analyses of human and murine AAA scRNA-seq datasets identified crosstalk between stromal and immune cells,^15–17^ implicating them in AAA pathogenesis. However, scRNA-seq analyses are significantly limited by low cell yield, biased cell recovery, and loss of spatial context to infer cell–cell communication. Spatial transcriptomics allows gene expression to be mapped in situ—enabling precise localization of disease-driving cells and their crosstalk in intact tissue.^18,19^ Despite their marked regional heterogeneity, AAAs have not been systematically analyzed with spatial transcriptomics.

*CXCR4* is a potential marker of inflammatory activity in the AAA wall,^21^ and the *CXCL12–CXCR4* axis regulates immune cell recruitment, progenitor cell trafficking, and tertiary lymphoid structure formation in vascular disease^22,23^ and drives profibrotic programs in cardiac fibrosis.^24^ This axis has not been analyzed in human AAA wall using spatial transcriptomics. In this study, we used the Xenium spatial transcriptomics platform, which provides subcellular spatial resolution of 5001 RNA targets, allowing near single-cell readouts^20^, to create a high-resolution spatial transcriptomic atlas of human AAA. By integrating transcriptomic profiling and histological analysis of AAA specimens, we found increased crosstalk between fibroblasts expressing *CXCL12* and immune cells expressing *CXCR4*. Our findings show how spatial biology can be applied to translational research in vascular disease and identify the *CXCL12–CXCR4* axis as a potential therapeutic target for AAA.

## METHODS

### Study Approval

The study was approved by the Institutional Review Board of the University of California, San Francisco (19-28098 and 20-31518) and conducted according to the Declaration of Helsinki principles

### Study Participants and Biospecimen Collection

AAA samples were prospectively collected from patients ≥18 years-old undergoing open AAA repair. Patients with mixed connective tissue disorders, active infection, chronic liver disease (Child-Pugh ≥B), stage-5 chronic kidney disease or creatinine ≥2 mg/dL, chronic inflammatory disorders, or major surgery or illness requiring hospitalization within 30 days were excluded. Consent was obtained from all patients before the surgery. Clinical data were retrieved from medical records.

Control samples were obtained from organ donors ≥18 years old without a history of aortic disease, human immunodeficiency virus infection, atherosclerotic disease, vasculitis, chronic hepatitis, or autoimmune diseases. All controls had given consent for use of their organs for research. Corresponding de-identified clinical data were obtained.

Samples of full-thickness AAA and control infrarenal abdominal aorta (1 x 4 cm) were excised and immediately placed into chilled phosphate-buffered saline, fixed in 10% formalin for 24–48 hours, washed with phosphate-buffered saline, placed in 70% ethanol for up to a month, and embedded in paraffin.

### Tissue Preparation

Tissue microarray blocks were constructed from the fixed tissues, with up to five 1-cm wide strips of full-thickness aorta sample embedded per block. For histological analysis, the blocks were cut into 5-µm sections, deparaffinized, rehydrated, and sequentially stained with Mayer’s hematoxylin and eosin (VWR 95057-844, 95057-848). Adjacent sections were stained with Verhoeff–Van Gieson stain (KTVELLT, Statlab) to identify elastin fibers. Elastin integrity was scored semiquantitatively (0 = intact, 4 = complete loss) as described.^25^

For spatial transcriptomic analysis, the blocks were cut into 5-µm sections, floated on an RNase-free 50°C water bath, mounted onto Xenium slides, air-dried for 30 minutes, baked at 65°C for 3 hours, placed in a desiccator at room temperature for 1–4 days, and processed with the manufacturer’s protocol (CG000578-Rev C, 10X Genomics) using the Xenium Human Gene Expression 5K panel, containing probes for 5001 genes (Supplementary Table 1). Slides were counterstained with DAPI, interior RNA stain, interior protein stain, and a membrane stain using the 10x Genomics Cell Segmentation Kit. Imaging and chemistry were done with the Xenium Analyzer software v3.2 according to the Analyzer User Guide (CG000584, Rev B, 10x Genomics). The run-output summary (*cells.csv.gz*), cell-feature matrix (*cell_feature_matrix.h5*), and transcript (*transcripts.csv*) files were used for analysis.

For immunofluorescent staining, 5-µm sections were baked at 65°C for 45 minutes, deparaffinized and rehydrated. Antigen retrieval was performed using an EDTA-based solution (Cell Marque #920P-04) and heated under pressure for 15 minutes. Slides were cooled, washed using PBS and PBS containing 0.1% Tween 20 (PBST) (Thermo Scientific #J20605-AP), blocked with 10% goat serum (Biolegend #927503) for 1 hour at room temperature (RT), then incubated with primary antibodies overnight at 4°C: PDGFRα (Biolegend #135901 1:200) and CXCL12/SDF-1 (R&D Systems #MAB350 1:200). Slides were washed with PBS and PBST, incubated with secondary antibodies (Invitrogen) for 1 hour at RT, washed with PBS and PBST, and mounted with ProLong Gold Antifade Reagent with DAPI (Thermo Scientific #P36931). Slides were imaged on a Nikon Crest LFOV Spinning Disk/C2 Confocal wide-field microscope and images analyzed using FIJI software.

### Data Preprocessing and Annotation

Spatial transcriptomic data were preprocessed with the Python packages scanpy,^26^ squidpy,^27^ and anndata.^28^ Cellular gene expression, metadata, and spatial location information were used to create anndata spatial objects for each sample. Low-quality transcripts (Xenium Q-score <20) were automatically excluded. Cells with <100 unique genes and genes detected in <10 cells were filtered out. scVI was used for batch correction by individual sample, and a principal component analysis with a latent space of 25 components^29^ was done. A neighborhood graph computed with n_neighbors=75 was embedded by using uniform manifold approximation and projection (UMAP)^30^ and clustered with the Leiden community detection method.^31^ The data were normalized and log-transformed, and the top 100 differentially expressed genes (DEGs) were analyzed by cluster to manually annotate each cluster using known markers. The top 5 DEGs were plotted by cluster.

### Unique Gene Threshold

In scRNA-seq analyses, cells with fewer than ∼200 detected genes are typically excluded to remove empty droplets and low-quality captures.^32^ We used a more permissive cutoff of 100 detected genes per cell to reflect differences between droplet-based scRNA-seq and spatial transcriptomics, which profiles a targeted panel of 5001 genes and begins with morphological segmentation of cells. Thus, the threshold primarily selects for technical quality rather than defining cell identity. Prior studies have used a threshold of 50–100 genes to retain low-transcript cell types such as neutrophils or neurons.^32–34^

To assess potential bias, we compared thresholds of 100 vs 50 genes. Lowering the cutoff added 199,249 cells (25.5% increase) and produced less distinct UMAP clustering. Two clusters were enriched in these cells (odds ratio [OR] ≥2, false-discovery rate [FDR] <0.05): a neutrophil cluster (cluster 18, 65% of cells with <100 genes, OR 5.47) and a *SOX2-OT* lncRNA/embryonic-like cluster with no discernable lineage markers (cluster 23, 76% of cells with <100 genes, OR 9.53), likely low-quality cells or cells with aberrant transcripts; using a threshold of 100, excluding these cells, was appropriate, resulting in higher quality of included cells and no overly biased exclusion of specific cell types (Supplementary Table 2).

### AAA vs Control Analysis

The cellular composition of each sample type (control and AAA) was analyzed. Differences were modeled with a beta-binomial regression, and the effect of cell type on disease was reported as an OR with 95% confidence interval. The results were compared to published scRNA-seq data^17^, including cell clusters identified, cells per sample, and overall distribution. In addition, each tissue sample was divided into “intima/media” or “adventitia”, and differences in cellular composition between AAA and control in each category were analyzed.

### Neighborhood Enrichment

Co-occurrence of cell clusters within a 50 µm radius was quantified with Squidpy’s neighborhood-enrichment analysis^27^ using Delaunay triangulation. Enrichment z-scores were calculated by comparing intercluster neighbor counts against a permutation-derived null, yielding a symmetric matrix of scores.

### Cell–Cell Communication

Intercellular communication was inferred separately in AAA and control samples with CellChat and CellPhoneDB.^35–38^ For each condition, all cluster pairs (source cell and target cell) were analyzed; ≥10 interactions and ligand and receptor expression in ≥10% of cells within their respective clusters was required for analysis. LIANA estimated permutation-based *P* values for each ligand–receptor cluster pair by shuffling cell labels within clusters, using raw counts and cell-type annotations. Significant interactions (*P*<0.05) were retained.

The plausibility of inferred interactions was assessed with a custom spatial score. For each ligand–receptor pair, the number of observed interactions within a 50-µm radius was calculated and then normalized by the sum of ligand^+^ source cells and receptor^+^ target cells. A 50-µm threshold was chosen to approximate the effective range of secreted factors in tissue.^39,40^ This approach incorporates local interaction density and adjusts for ligand/receptor expression outside spatially plausible contact zones.

### Immune Cell Microenvironments of CXCL12⁺ and CXCL12^−^ Fibroblasts

To compare immune landscapes, we analyzed the number and type of immune cells within a 50-µm radius of each *CXCL12⁺* and *CXCL12⁻* fibroblast in AAA samples. Differences in the number of immune subsets were determined with two-sided Fisher’s exact tests and the Benjamini–Hochberg correction (FDR<0.05); differences in the types of immune cells were determined with the Mann-Whitney U test (*P*<0.05). To assess the effects of distance, this analysis was repeated across radii from 25–400 µm.

### Genomic Analyses

To link AAA heritability to cell types, summary statistics from a genome-wide association study (GWAS) of AAA from the AAAgen Consortium, harmonized to GRCh37/hg19,^41^ were analyzed in relation to our data. Gene-based associations were tested with MAGMA (Multi-Marker Analysis of GenoMic Annotation (v1.10), the default single-nucleotide-polymorphism–wise mean model, and the 1000 Genomes Phase 3 European reference panel.^42^ Variants not in the reference or in ≥50 cells were excluded. For gene-property analyses, we tested continuous cell-type properties derived from the sets of DEGs (log fold-change >0 and *P*<0.05) used to define our cell clusters (see Supplementary Table 3). Models were fit as linear regressions of MAGMA gene Z-scores on each property while conditioning on average gene expression. FDR<0.05 was considered significant. Differential expression of AAAgen Consortium genes in our clusters was also analyzed.

## RESULTS

Infrarenal abdominal aorta tissue samples were collected from 11 patients undergoing open AAA repair and 12 deceased brain-dead donors (controls) undergoing organ harvest at the University of California, San Francisco between October 2023 and May 2025. Their mean ages were 72±6 and 53±13 years, respectively (Table 1). Most AAA patients had a history of smoking and similar comorbid conditions; their average maximal aortic diameter was 6.3±1.4 cm.

**Table 1.**
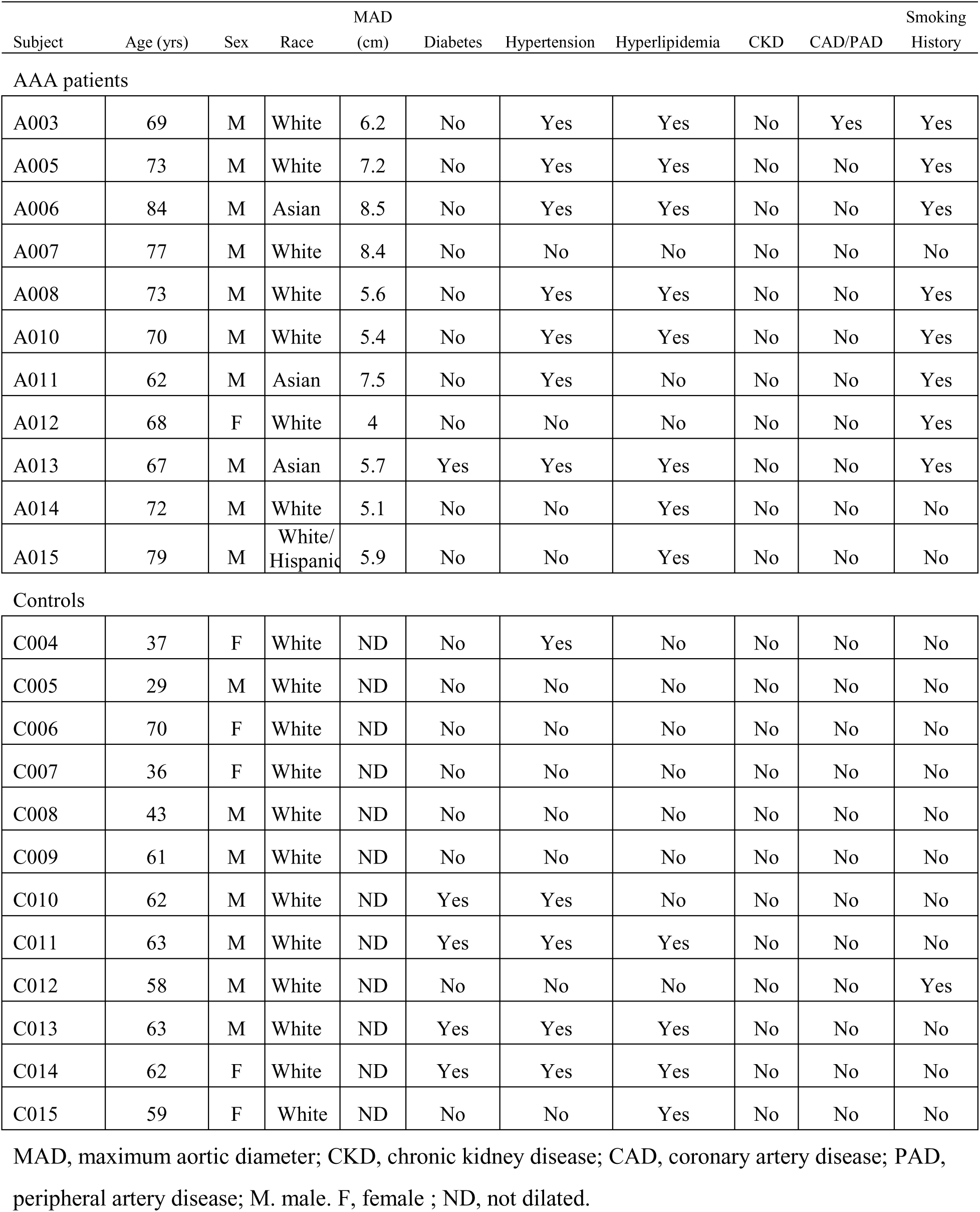
Patient demographics, medical history, and imaging findings.

### AAAs Have Distinct Histopathological and Structural Features

Control aortas appeared healthy. The intima was intact, the media had a thick band of SMCs, and the adventitia was thin and lacked clear evidence of adventitial tertiary lymphoid organs (ATLOs), inflammatory infiltrates, and atherosclerotic degeneration (Figure 1A). All AAAs showed intimal and medial degeneration, loss of medial SMCs, adventitial fibrosis, high-density cellular infiltrate (likely inflammatory and most notable in the adventitia and outer media), and neovascularization, and some had atherosclerotic plaque and calcification (Figure 1B–D), consistent with previous reports.^43^ The dense cellular infiltrates at the medial/adventitial border and throughout the adventitia were consistent with the histological makeup of ATLOs (Figure 1C)—organized immune hubs of T cells, B cells, plasma cells, and lymphatics/high endothelial venules that contribute to chronic inflammation, matrix remodeling, and structural weakening in AAA.

**Figure 1.**
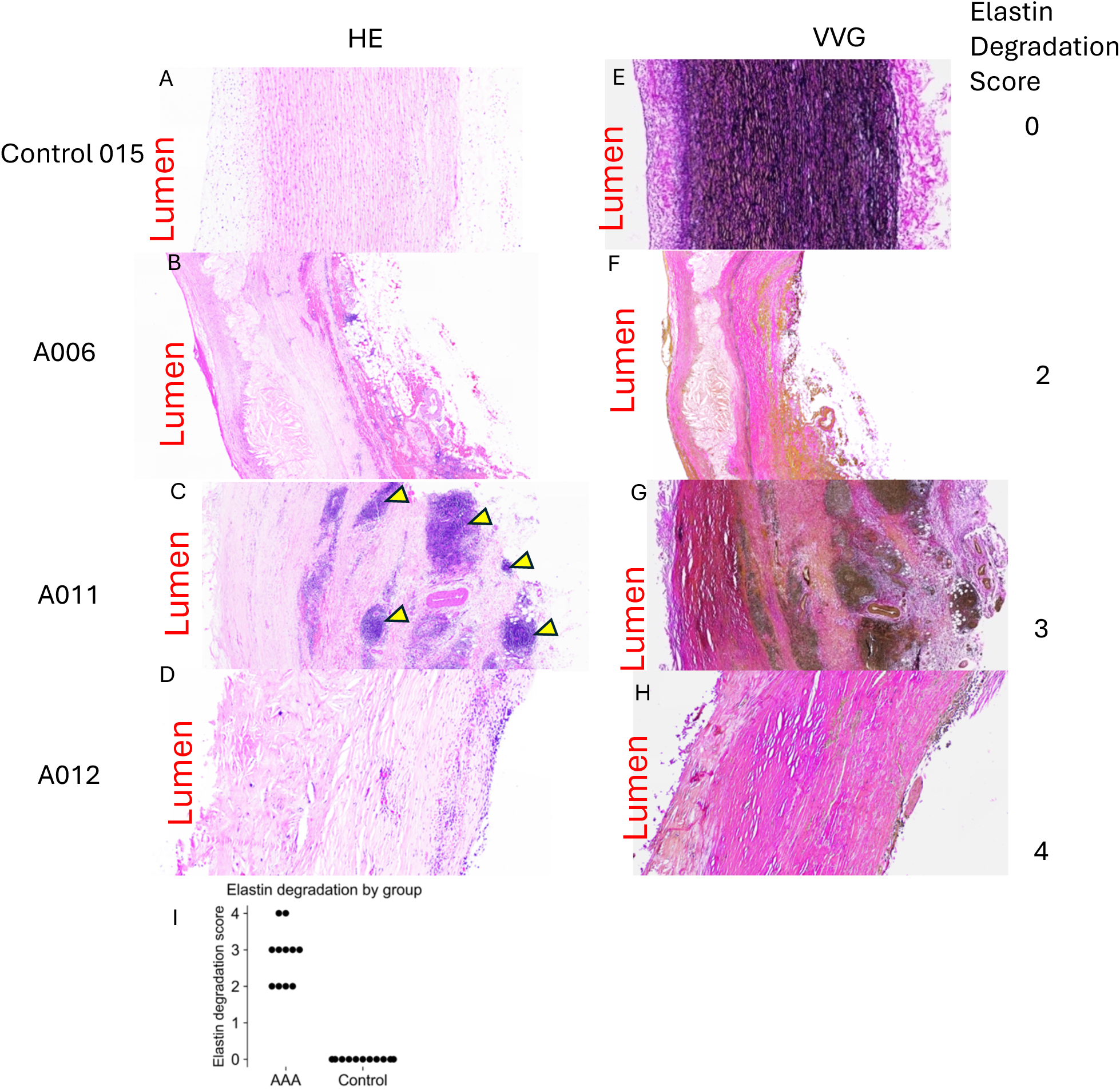
Paired sections (5-μm) of a control aorta and of AAA samples from three patients and corresponding elastin degradation scores. Sections were stained with hematoxylin/eosin (A–D) and Verhoeff-Van Gieson stain (E–H). Original magnification, 10×. **A,** Control aorta with thin adventitia, compact, regular, intact media with thick band of homogeneous cells, and loose adventitial tissue with sparse nuclei. **B–**D, AAA samples showing atherosclerotic plaque, disrupted/thin, and dense cellular infiltrate most notable in the adventitia(**B**); disrupted intima and dense cellular infiltrate (likely adventitial tertiary lymphoid organs) (arrowheads) in the outer media and through the adventitia (**C**); and disrupted intima with surrounding atherosclerotic changes, intimal and medial degeneration, and adventitial cell infiltration (D). **E,** Control aorta showing regular, intact elastic lamellae throughout the media. **F–H,** AAAs showing thin elastic laminae with moderate elastin fragmentation (**F**); severe fragmentation and disruption of elastin fibers (**G**); and near-complete loss of elastin (**H**). **I,** Elastin degradation scores by group.

Control aortas had regular, intact elastic lamellae without breaks, fraying, or loss of elastin (Figure 1E). AAA samples had elastin degeneration, a hallmark of AAA,^44^ ranging from thin elastic lamellae to no lamellae (Figure 1F, G, I). The elastin integrity score was 0 in control aortas, indicating no degeneration, and 2–4 in AAAs (Figure 1I), indicating vascular remodeling and advanced disease.^25^

### Spatial Transcriptomics Reveal a Diverse Atlas of Stromal and Immune Cells

To identify cellular and molecular changes, we compared the spatial transcriptomics^45^ of all AAA and control samples. Cell staining and segmentation identified 147,182 control and 434,482 AAA cells (Figure 2A–H), with an average of 324 transcripts and 241 genes per cell (Figure 2G, Supplementary Figure 1). Unsupervised clustering of dimensionally reduced transcriptional profiles identified 26 clusters. Differential gene analysis (Supplementary Figure 2) was used to annotate vascular and immune cells based on scRNA-seq datasets (Supplementary Figure 3).^17,46–48^ Twelve cell types were identified: SMCs, B cells, macrophages, T cells, endothelial cells (ECs), fibroblasts, monocytes, dendritic cells, neural cells, neutrophils, mast cells, and adipocytes (Figure 2G, Supplementary Table 3). Stromal cells accounted for 90% of cells in control aortas but only 32% in AAA, reflecting a marked immune expansion (66.9% vs 10% in controls) (Figure 2H, 2I, Supplementary Table 4). Neighborhood enrichment analysis was performed to determine propensity for spatial proximity among cell clusters (Figure 3A), and tissues were divided into “intima/media” and “adventitia” to analyze differences across the aortic wall (Figures 3B-C).

**Figure 2.**
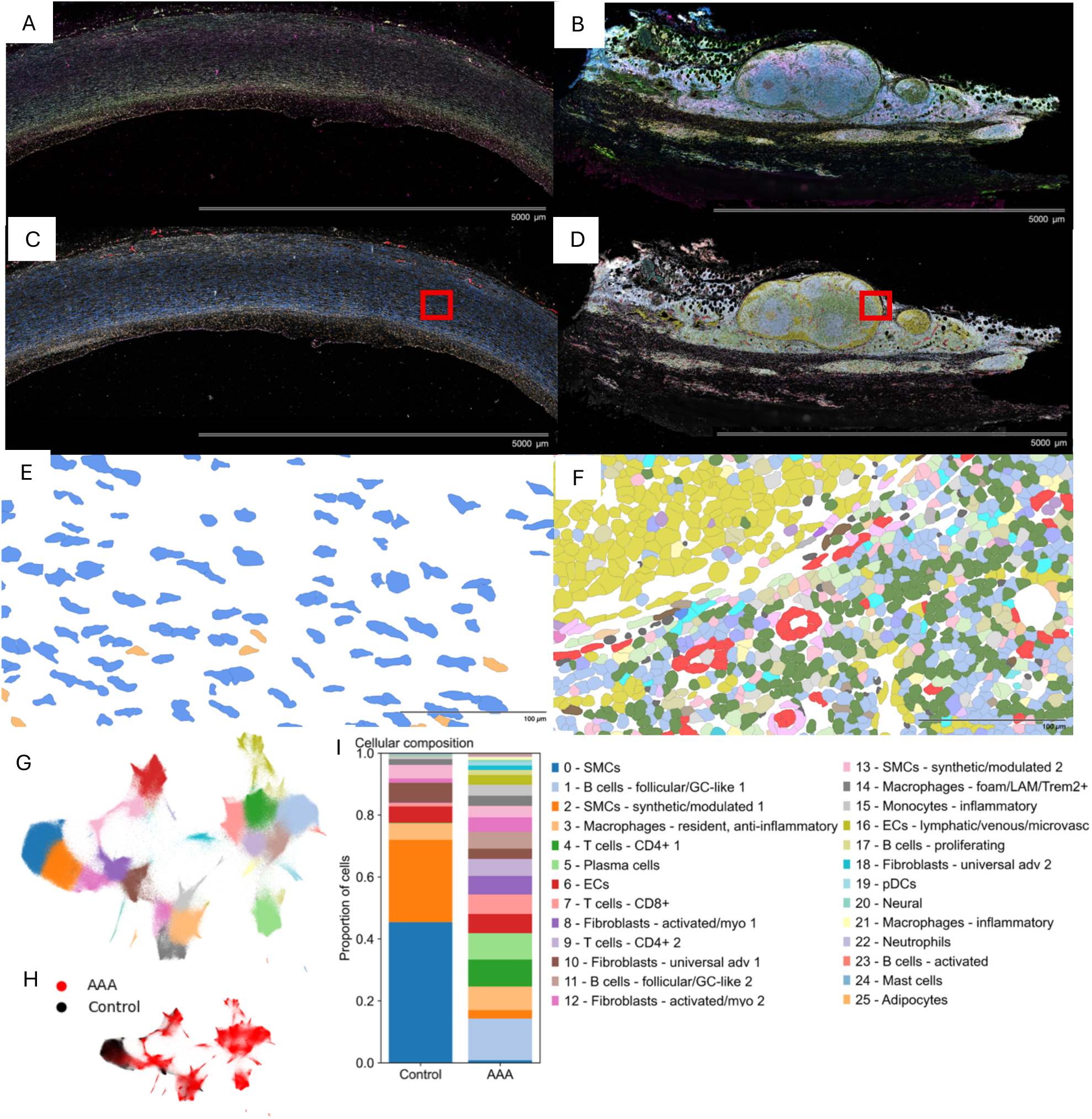
Xenium spatial transcriptomic cell segmentation and clustering, with comparison between control and AAA. Sections (5 μm) of control aorta (**A**) and abdominal aortic aneurysm (AAA) (**B**) stained with DAPI, interior RNA, interior protein, and cell-membrane stain and corresponding cell segmentation grayscale overlay of the stained control (**C**) and AAA (**D**) sections. Original magnification 40×. Higher magnification of the areas indicated by red rectangles in **C** (**E**), and **D** (**F**). **G,** Unbiased Leiden clustering of batch-corrected, dimensionally reduced transcriptional profiles from 581,664 cells isolated from 12 control aortas and 11 AAA identified 23 cell lineages in 26 clusters. **H,** UMAP by sample type. AAA cells (n=434,482) were found mostly in immune clusters, while control aorta cells (n=147,182) were mostly in stromal clusters. pDCs, plasmacytoid dendritic cells. **I**, Overall cellular composition by sample type.

**Figure 3.**
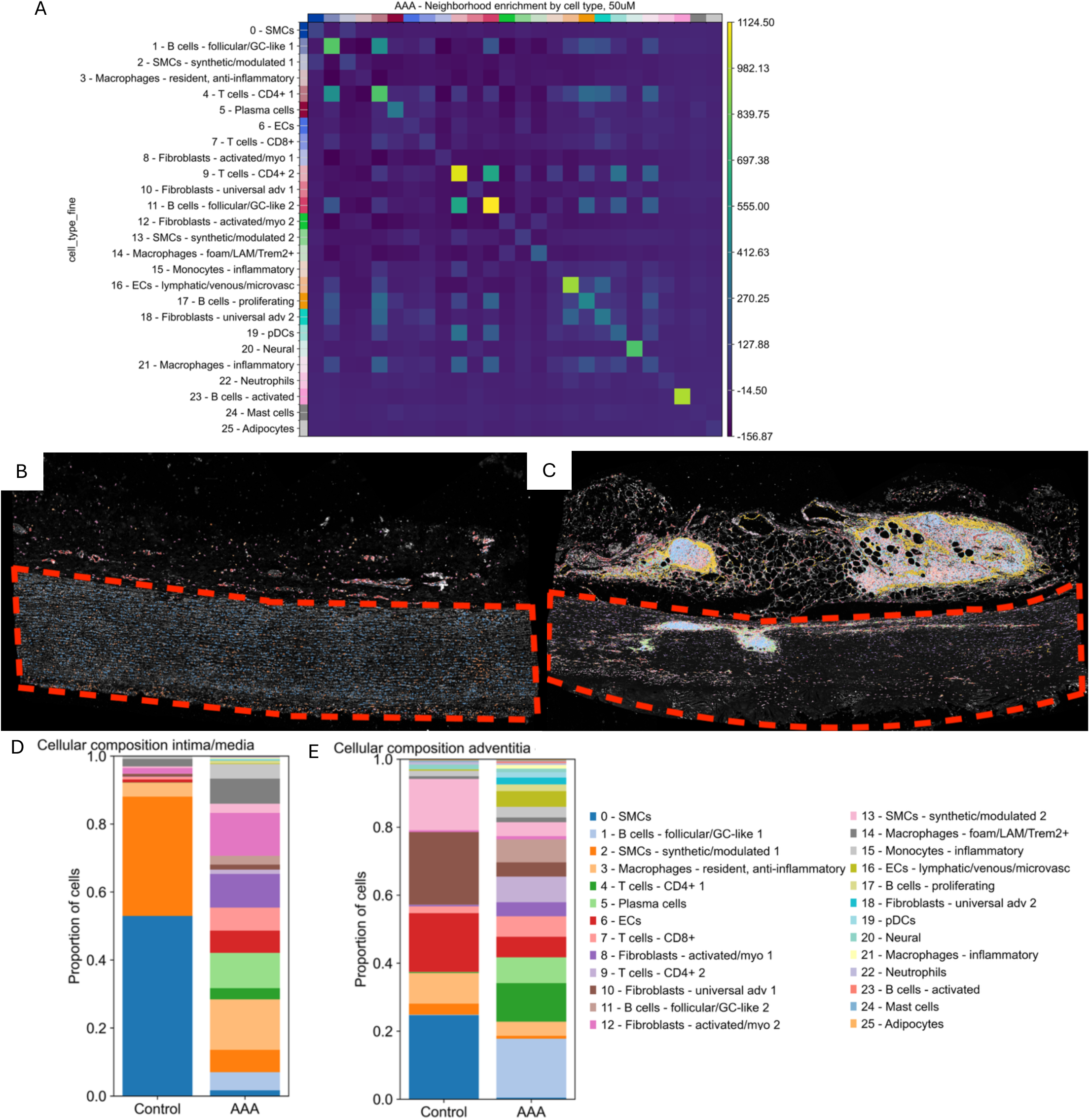
Spatial analyses reveal cells likely to co-localize in abdominal aortic aneurysm (AAA), and cellular composition differences across the aortic wall between control and AAAA. Neighborhood enrichment plot for AAA samples, indicating cell types most likely to colocalize within a 50-μm radius of one another. **B** and **C**, Spatially resolved clusters with grayscale segmentation staining overlay on sections of control aorta (B) and AAA (C), oriented with the adventitia on top. Red outline indicates intima/media. **D** and **E**, Cellular composition of intima/media (**D**) and adventitia (**E**) in control aorta and AAAs.

### SMC Populations

Healthy aortas were primarily composed of two subpopulations of SMCs^49^ (71% of cells) in the intima and media (Figures 3D-E). Cluster 0 (45% of control cells), consisted of contractile SMCs expressing *MYH10,*^50^ *SMTN,*^51^ and *RGS5*^52^ (Figures 2G, 2I, Supplementary Tables 3–4). This cluster was almost completely depleted in AAAs (0.8%) (Supplementary Table 4), consistent with previous reports^53^ and possibly due to apoptosis driven by cytotoxic mediators secreted by infiltrating immune cells.^54^ Cluster 2 (26.6% of control cells) consisted of synthetic/modulated SMCs that expressed the SMC genes *MYH9/10,*^50^ *ACTN1/4,*^55^ and *MYLK,*^56^ the fibroblast-associated genes *PDGFRα,*^57^ *CCN1/2/3/5,*^58^ and *COL4A1/2,*^59^ and the elastic fiber remodeling genes *LTBP1/2,*^60^ *LOXL1,*^61^ and *TNFRSF11B*^62^ (Figures 2G, 2I, Supplementary Tables 3-4) and resembled fibromyocytes.^63,64^ In AAAs, these cells were nearly absent (2.7%) (Supplementary Table 4).

Cluster 13 (synthetic/modulated SMCs similar to those in Cluster 2) was rare in controls (4.4% of cells) but was the predominant SMC population in AAAs (3.6%) (Figure 2I, Supplementary Table 4). These cells expressed the SMC-related genes *NOTCH3,*^65^ *RBPMS,*^66^ and *MYLK,*^56^ the fibroblast/ECM remodeling genes *COL4A1/2,*^59^ *CCN1*^58^*, HSPG2*^67^, and *AEBP1*,^68^ and the stress–angiogenic/vasoactive markers *EPAS1,*^69^ *ADAMTS1/4,*^70,71^ and *THBS1*^72^ (Figure 2G, Supplementary Table 3). These genes have been implicated in hypoxia-driven SMC remodeling, ECM degradation, angiogenesis, and altered vascular tone,^71,73–75^ consistent with phenotypic modulation of SMCs in AAA toward a pro-inflammatory synthetic state that drives pathological remodeling.^76^ Cluster 13 synthetic SMCs were predominantly located in the adventitia of control aortas and AAAs (Figure 3D and E).^77^

These findings suggest two SMC phenotypes in the aortic wall: one skewed toward matrix repair and calcification and more prominent in healthy aorta and one skewed toward hypoxia adaptation, angiogenesis, and altered vascular tone in AAA.

### Fibroblast-Like Cells

Fibroblasts maintain vascular tissue integrity^78^ but can drive maladaptive remodeling under pathological conditions.^79^ In AAA, they promote collagen turnover, elastin degradation, and secretion of proteases and cytokines, which weaken the aortic wall and promote adventitial neovascularization and immune cell recruitment.^80,81^

We found that fibroblasts were less abundant in control aortas than in AAAs (7.8% vs 15.5% of cells) (Supplementary Table 4). There were four subpopulations. Cluster 10 (universal adventitial fibroblast 1), the largest fibroblast cluster in controls (3.3% of cells), located predominantly in the adventitia, expressed *PI16,*^82,83^ *PDGFRα,*^57^ *CCDC80,*^83^ *DPT,*^84^ and *CXCL12*^85^ (Supplementary Tables 3 and 4). In AAA, these fibroblasts comprised 6.3% of cells and were also located in the adventitia (Supplementary Table 4, Figure 3D and E). Cluster 18 (universal adventitial fibroblast 2), located predominantly in the adventitia (Figure 3D and E), also expressed those genes, along with *CXCL12*, *TNC,*^86^ *DPT,*^84^ and *SERPINE1*^87^ (Figure 2G, Supplementary Table 3). Cluster 18 represented 1.4% of cells in AAAs but only 0.0007% of cells in control aortas. These clusters resemble two universal fibroblast archetypes: PI16⁺ adventitial fibroblasts (structural, progenitor-like), which are quiescent in uninjured tissues,^88^ and *CXCL12⁺* stromal niche fibroblasts (immune-interacting and paracrine hubs),^84^ which colocalize with lymphocytes and inflammatory macrophage (Figure 3A) and likely help shape the tissue microenvironment through *CXCL12* signaling to immune cells.

Two populations of activated fibroblast/myofibroblast present predominantly in the intima/media (Figure 3D and E) were less abundant in control aortas (1.5% vs. 10.7% of AAA cells) (Supplementary Table 4). Cluster 8 (activated fibroblast/myofibroblast 1), the largest fibroblast cluster, was enriched in cells expressing the ECM-remodeling and matricellular genes *THBS1/2,*^72,89^ *COMP,*^90^ *COL5A1/2,*^91^ *CCN2,*^58^ *SERPINE1,*^87^ *SMOC2,*^92^ *ADAMTS2,*^71^ *CTHRC1,*^93^ and *MMP14.*^94^ Cluster 12 (activated fibroblast/myofibroblast 2) had a similar profile but expressed high levels of *POSTN*^95^ (Figure 2G, Supplementary Table 3). These populations, defining a *POSTN/THBS*-high subset with wound-healing and fibrotic properties, closely resemble the fibroblast-Cilp (*Postn*^+^/*Adamst2*^+^) and fibroblast-Thbs4 (*Smoc2*^+^/*Cthrc1*^+^/*Comp*^+^) subsets that contribute to pathological fibrotic remodeling of cardiac ECM and cardiac fibrosis in mice,^96^ and *POSTN*^+^ fibroblasts, which are highly proliferative, express contractile markers, and may contribute to cardiac fibrosis after myocardial infarction.^97^

These findings define two fibroblast axes in AAA: *PI16^+^**/**CXCL12⁺* adventitial fibroblasts that match conserved archetypes and POSTN/THBS-high activated myofibroblasts also seen in cardiac fibrosis.

### ECs

scRNA-seq has limited ability to detect ECs in vascular preparations after tissue digestion.^98,99^ Since luminal ECs were likely lost during intraluminal thrombus removal before AAA excision, the two endothelial subsets we identified were primarily in the adventitia (Figure 3D and E). Cluster 6 ECs were enriched in stress-adapted, pro-angiogenic genes with classical endothelial markers (*PECAM1,*^100^ *CD34*^101^). Cluster 16 (lymphatic/microvascular ECs) exhibited hypoxia-driven activation, expressed permeability and proliferation markers *(EPAS1*^102^*, PLVAP*^103^), had a lymphatic/venous microvascular profile (*FLT4,*^104,105^ *CAVIN2*^106^), and co-expressed *ACKR3,* a CXCL12 receptor that promotes atherosclerosis by increasing arterial adhesion and immune cell infiltration^107^ (Figure 2G, Supplementary Table 3). Identified in both control (5.6%) and AAA tissues (6.2%) (Supplementary Table 4), Cluster 6 ECs colocalized with CXCL12⁺ universal adventitial fibroblast cluster 2 (Cluster 18) and synthetic/modulated SMCs (Clusters 2 and 13) (Figure 3A), likely enabling greater recruitment of inflammatory cells by increasing neovascular permeability. Cluster 16 ECs, present almost exclusively in AAA adventitia (3.2% vs 0.1% of control) (Supplementary Table 4), colocalized with Cluster 18 and lymphocytes (Figure 3A), consistent with lymphangiogenesis and angiogenesis in hypoxic and inflamed areas of the aortic wall, a feature of end-stage AAA.^108,109,110^

### Myeloid Cells Accumulate in AAAs

We identified three macrophage clusters. Cluster 3, resident anti-inflammatory macrophages expressing homeostatic genes (*STAB1,*^111^ *CD163,*^112^ *MRC1*^113^), was the dominant immune cluster in control tissues; it was present throughout the wall and preferentially in the intima/media in AAA. Cluster 14, foamy, lipid-associated TREM2^+^ macrophages expressing lipid-handling^114^ and lysosomal genes (*TREM2,*^75,102^ *APOE,*^91,114^ *LPL*^91,114^) (Figure 2G, Supplementary Table 4) that are pro-inflammatory and pro-angiogenic in AAAs,^16^ was mostly identified in intima/media in AAAs and control aortas. Cluster 21 (inflammatory) expressed *MMP9,*^115^ *CTSC,*^116^ *CTSL,*^117^ and cytokine/chemokine genes, consistent with matrix-degrading, pro-inflammatory activity, was identified only in AAA and was found throughout the aortic wall. Cluster 15, inflammatory monocytes expressing *CD14*^118^, *S100A8/9*^119,120^, and *IL1B,*^121^ were abundant throughout the aneurysmal and healthy aortic wall and resembled a dendritic-like subset found to contribute to tissue injury through innate immune activation, cytokine amplification, and lymphocyte recruitment.^120^ Cluster 19, plasmacytoid dendritic cells expressing *IRF7* and *CLEC4C*, were rare and found only in AAA. Cluster 22, neutrophils expressing *FPR1, CXCR1,* and *CSF3R,* and Cluster 24, mast cells expressing *KIT* were slightly more prevalent in AAAs (Figure 2G, Supplementary Table 4).

### B Cells

Lymphocytes were rare in control aortas (1.4% of cells) but made up 49.8% of total cells and >74% of all immune cell infiltrate in AAAs (Supplementary Table 4). In contrast to flow cytometry and scRNA-seq results suggesting that T cells are the predominant lymphocytes in AAAs,^17,124,125^ we found that in 8 of the 11 AAA samples (Supplementary Figure 4A–C), T cells were less abundant than B cells (20.5% vs 29.3% of all AAA cells). B cells likely sustain immune activation via antibody and cytokine production within ATLOs, promoting aortic wall weakening and aneurysm progression.^110^

We identified five B-cell populations, mostly in the adventitia, that preferentially colocalized with T cells (Figure 3A, D and E), consistent with local B-cell maturation and antibody secretion in ATLO niches. Cluster 1 (B cells) was the most abundant B-cell population in AAAs (13.4%) but rare in healthy aortas (0.05%) (Supplementary Table 4). These cells were rich in canonical B-cell markers and markers of B-cell receptor signaling and survival and resembled mature follicular/germinal center-like cells (MS4A1,^114^ CD19,^114^ CD79A,^114^ *TNFRSF13C,*^115^ *BCL2,*^116^ *CXCR4*^117^). Cluster 11 (B cells) accounted for 5.4% of cells in AAAs and was enriched in similar markers but expressed higher levels of *CXCL13,*^118^ *CR2,*^119^ and *CD83*^119^ and lower levels of *CXCR4.*^117^ Cluster 5 (plasma cells expressing *XBP1,*^120^ *MZB1,*^121^ and *PRDM1*^122^) was almost absent in control aorta (0.05%) but accounted for 8.5% of cells in AAA (Supplementary Table 4). Cluster 17 (proliferating B-cells expressing TUBB^123^ and STMN1^123^) was located in areas of dense immune infiltrate in the aneurysmal adventitia (Figure 3A and E), likely indicating active germinal center expansion. Cluster 23, activated B cells expressing CD83^124^ (Figure 2G, Supplementary Table 4), which could increase immune activation and chronic inflammation, was found in AAA adventitia.

### T Cells

CD4 and CD8 T cells are thought to clonally expand in AAA.^128^ CD4 T cells promote AAA progression by sustaining chronic inflammation, inducing macrophage polarization and SMC dysfunction through cytokines, and CD8 T cells promote aortic wall destruction through IFN-γ-driven apoptosis and matrix metalloproteinase activation.^125–127^ We identified two clusters of CD4 T cells and one cluster of CD8 T cells. Cluster 4 CD4 T cells were enriched in memory, activation, and T-regulatory markers (CD4, *SELL,*^111^ CTLA4,^112^ and TGIT^112^) (Figure 2G, Supplementary Table 3) and located in areas of dense adventitial infiltrate (Figure 3D and E). They were the most abundant T cells in AAAs (8.7%) but rare (0.09%) in control aortas (Supplementary Table 4). Cluster 9 CD4 T cells, which resembled Cluster 4 T cells and localized to the adventitia but were not activated (Figure 2G, Supplementary Table 3), were abundant in AAAs (5.7%) but nearly absent in control aorta (0.03%) (Supplementary Table 4). Cluster 7 *CD8* T cells were enriched in cytotoxic markers (CD8A, *GZMB,*^113^ *PRF1*^113^) (Figure 2G, Supplementary Table 3). Present throughout the aneurysmal wall and healthy adventitia, they constituted 6.3% of all AAA cells and were the predominant T cell in control aortas (1.1% of cells) (Figure 3D and E, Supplementary Table 4).

### Fibroblast-Driven Immune Signaling Is Increased in AAAs

Next, we used CellChat and CellPhoneDB^129^ to examine intercellular communication networks and a custom spatial scoring metric to assess the plausibility of ligand–receptor interactions within a 50-µm radius. In control aortas, these interactions were fewer and of lesser magnitude than in AAAs (Figure 4A and B) and largely centered on homeostatic signaling such as canonical Notch signaling (*JAG1-NOTCH1/2/4*)^65,130^ and efferocytosis and anti-inflammatory signaling (*GAS6/AXL, GAS6/MERTK)*^131,132^ (Figure 4A). In contrast, AAA tissue displayed a marked expansion of fibroblast-driven interactions, particularly *CXCL12–CXCR4* signaling from fibroblasts to lymphocytes (Figure 4B) and *SEMA4D-PLXNB2* signaling from leukocytes to stromal cells and macrophages (Figure 4B), which is associated with immune-driven angiogenesis, stromal activation, and inflammatory cell recruitment.^133,134^ Interactions between immune cells were also selectively enriched in AAA (Figure 4B). Importantly, the spatial score highlighted that the highest-ranking interactions were not only statistically significant but also spatially plausible. The *CXCL12–CXCR4* interaction between Cluster 18 (fibroblasts) and Cluster 4 (CD4 T cells) (Figure 4D) was particularly prominent in AAA adventitia and, to a lesser degree, in the outer media. The spatially agnostic CellChat scores did not always correlate with the spatial score; some interactions with similar CellChat scores had different spatial scores (Figure 4B and C), highlighting the need for spatial consideration in rating interactions. These findings highlight the potential role of paracrine signaling in aneurysm pathogenesis.

**Figure 4.**
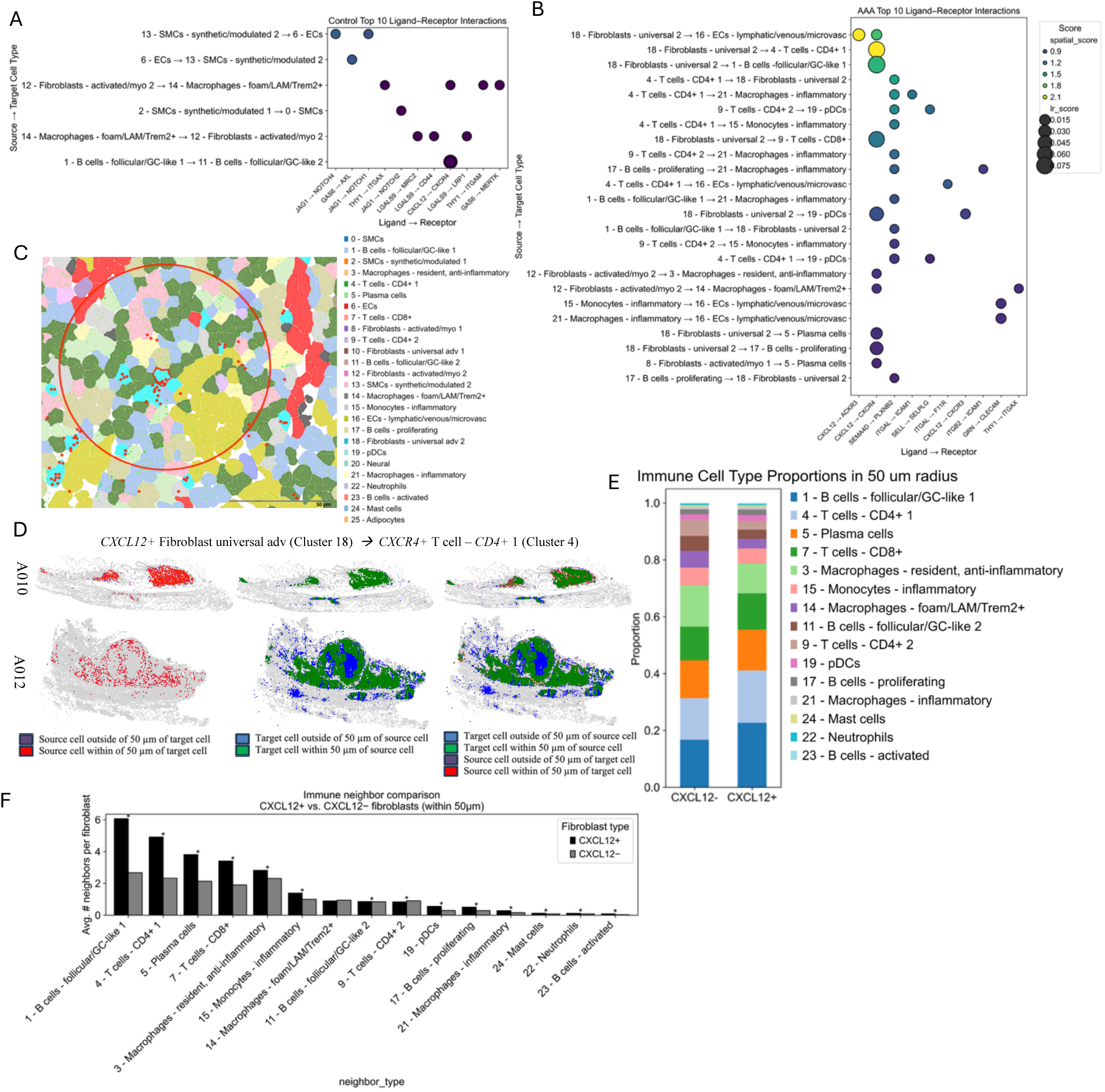
Cell-cell communication in control and abdominal aortic aneurysm (AAA) within a 50-µm radius and analysis of the immune microenvironment of CXCL12^+^ and CXCL12^-^fibroblasts. **A and B,** Top 10 spatially resolved ligand–receptor interactions in control aortas (**A**) and AAA (**B**) by spatial score. **C**, Cell segmentation plot showing CXCR4^+^ immune cells within 50-μm (red circle) of a CXCL12^+^ fibroblast (aqua cell outlined in red). Red dots represent CXCL12 transcripts; green dots represent CXCR4 transcripts. **D,** Spatial representation of signaling from CXCL12^+^ universal 2 fibroblasts to CXCR4^+^ CD4 T cells (one of the top spatial ligand-receptor interactions) in 2 AAA samples. Source cells within vs outside of 50µm of target cells are in the first column, vice versa in the second column, and a combined depiction in the third column. **E,** Proportions of immune cell types within 50 μM of CXCL12^+^ and CXCL12^−^ fibroblasts. pDCs, plasmacytoid dendritic cells. **F,** Average number of neighboring immune cell types within a 50-μm radius of CXCL12^+^ vs CXCL12^−^ fibroblasts. \**P*<0.05 by Mann Whitney test.

### CXCL12⁺ Fibroblasts Are Present in Immune-Cell-Rich Niches in AAAs

To dissect the immune microenvironment of *CXCL12*⁺ fibroblasts, we compared their spatial neighborhoods (within a 50-µm radius) to those of *CXCL12*⁻ fibroblasts. *CXCL12* was primarily expressed in fibroblasts and ECs (Supplementary Figure 5A). At a protein level, CXCL12 was also found to colocalize with PDGFRα, a fibroblast marker (Supplementary Figure 5B).

*CXCL12*⁺ and *CXCL12^−^* fibroblasts had similar immune cell neighbors (mostly lymphocytes and macrophages) (Figure 4F). *CXCL12⁺* neighbors were slightly enriched for follicular-like B cells (Cluster 1), *CD4* T cells (Cluster 4), plasma cells (Cluster 5), and *CD8* T cells (Cluster 7) and slightly depleted of resident and foam/TREM2⁺ macrophages (Cluster 14), follicular-like B cells (Cluster 11), *CD4* T cells (Cluster 9), and inflammatory monocytes (Supplementary Table 6). *CXCL12*⁺ fibroblasts had more immune cell neighbors per cell than *CXCL12*⁻ fibroblasts, most notably follicular-like B cells (Cluster 1), *CD4* T cells (Cluster 4), plasma cells (Cluster 5), resident macrophages (Cluster 3), and plasmacytoid dendritic cells (Cluster 19) (Figure 4G). Areas around *CXCL12⁺*fibroblasts were significantly enriched for all immune cells except foamy, lipid-associated *TREM2*^+^ macrophages (Cluster 14) (Figure 4G) and a *CD4* T-cell cluster (Cluster 9), which were slightly enriched around *CXCL12⁻* fibroblasts. The presence of *CXCL12*⁺ fibroblasts in lymphoid- and myeloid-rich niches aligns with their role as paracrine organizers of immune cell recruitment.

When neighborhood radius was expanded, *CXCL12*⁺ fibroblasts had more immune neighbors per cell than *CXCL12*^−^ fibroblasts, with similar composition across radii (Supplementary Figure 5C and 5D), indicating they serve as focal hubs for lymphoid clustering within broader immune-infiltrated zones.

**Figure 5.**
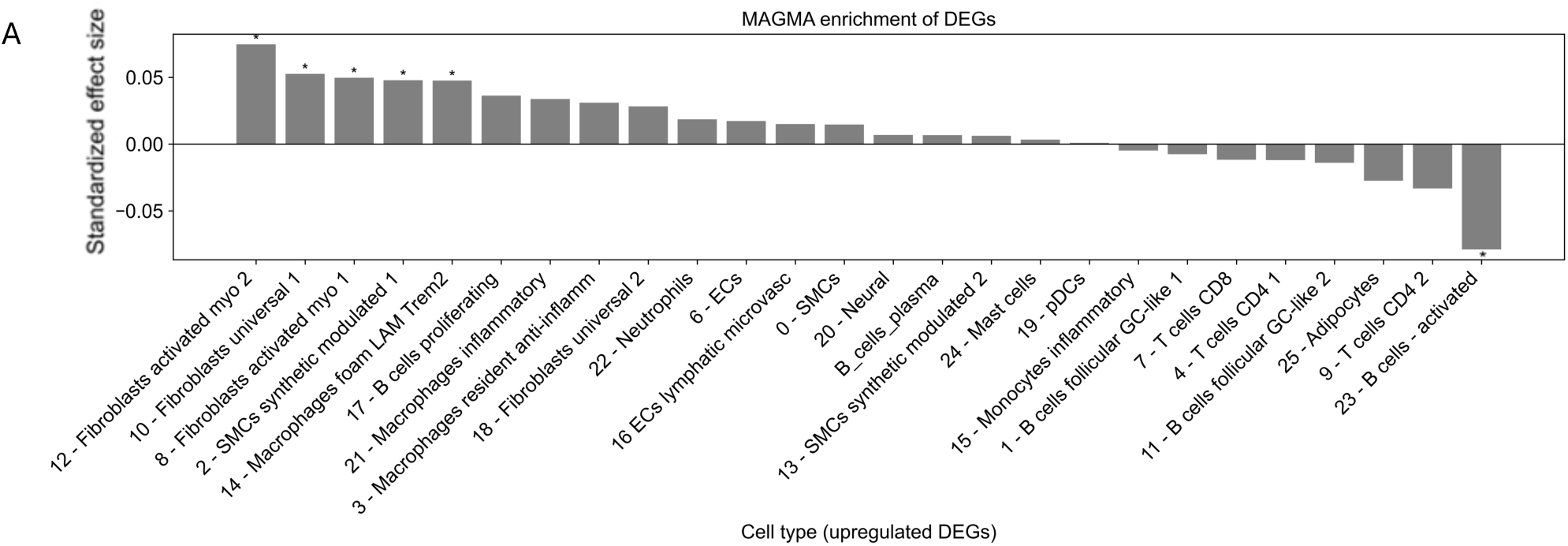
Multi-marker analysis of genomic annotation (MAGMA) analysis of enrichment vs depletion of genome-wide association study (GWAS) loci within clusters. Activated (myo)fibroblast clusters (Cluster 8 and 12) and universal fibroblast cluster 1 (Cluster 10) showed the strongest enrichment, surpassing FDR correction (*q<0.05), followed by synthetic/modulated SMCs and foam-like/Trem2⁺ macrophages. Adaptive immune cell clusters, including T cells and B cells, showed no significant enrichment, and activated B-cells (Cluster 23) was significantly depleted.

### AAA Genetic Risk Is Linked to Fibroblast, SMC, and Macrophage Clusters

To link transcriptional states to genetic risk, we analyzed GWAS data from the AAAgen Consortium^41^ with MAGMA.^42^ Regression of cluster-defining DEGs against AAA gene-level associations to test for enrichment in AAA loci identified cell types that may be responsible for genetic risk. Activated (myo)fibroblasts (Clusters 8 and 12) and universal fibroblasts 1 (Cluster 10) showed the strongest enrichment, followed by synthetic/modulated SMCs (Cluster 2) and foamy macrophages (Cluster 14) (Figure 5). Adaptive immune cells showed no enrichment, and activated B cells (Cluster 23) were depleted (Figure 5). Thus, stromal populations, particularly activated fibroblasts, may contribute disproportionately to AAA heritability, aligning with their transcriptional expansion in aneurysmal tissue and their role as paracrine organizers of immune microenvironments. We then mapped GWAS loci-mapped genes to cluster expression (Supplementary Figure 6). Activated (myo)fibroblasts (Cluster 8 and 12) and universal fibroblasts 1 (Cluster 10) expressed high levels of *LRP1, TGFBR2, MRC2, CDKN1A, COL4A2,* and *THBS2*, and foamy macrophages (Cluster 14), expressed high levels of *LRP1, MMP9, CDKN1A, LIPA,* and *PLAUR* (Supplementary Figure 6).

## DISCUSSION

Using state-of-the-art transcriptomics on FFPE samples we were able to develop the first-ever subcellular-resolution spatial transcriptomic atlas of human abdominal aortic aneurysm (AAA), with >580,000 cells identified from aortic tissue sections. This was made possible by recovering substantially more high-quality cells per patient than scRNA-seq studies,^17^ enabling us to map stromal and immune states that drive AAA and cell–cell communication in intact tissue at high resolution. In addition to all major clusters from prior work^17^ except NK cells, we found clusters of neural cells, adipocytes, mast cells, diverse B-cell lineages, and a higher proportion of stromal cells, especially SMCs. Immune cells were largely distributed as previously reported; however, we found relatively more B cells than T cells and fewer myeloid cells.

Our spatial analysis showed that *CXCL12*⁺ fibroblasts, mostly in the adventitia, are central organizers of adventitial immune niches. Cell–cell communication analysis revealed a significant interaction between these fibroblasts and *CXCR4⁺*immune cells. *CXCL12*⁺ fibroblasts were preferentially surrounded by primarily lymphocyte subsets and thus may act as stromal hubs that coordinate adaptive immune clustering and shape the adventitial microenvironment into ATLO-like structures, enabling chemokine-driven crosstalk, findings that scRNA-seq alone could not resolve. These fibroblasts along with myofibroblasts, an SMC cluster, and foamy macrophages were significantly enriched in AAA loci identified by GWAS, linking their transcriptional states to genetic risk. Thus, fibroblasts may orchestrate immune cell recruitment and positioning, linking stromal activation, immune cell recruitment, neovascularization and lymphoid-neogenesis in AAA and identifying stromal–immune crosstalk as a potential therapeutic target in AAA.

This finding is consistent with previous studies showing *CXCL12–CXCR4* signaling is upregulated in humans with AAA and in mouse models,^21^ and *CXCR4* depletion in mouse models suppresses AAA and decreases SMC apoptosis and phenotypic transformation, underscoring its functional importance in aneurysm biology.^137^ *CXCL12⁺* fibroblasts may increase microvascular permeability in cancer progression^138^ and in brain injury may create specialized stromal niches that retain T cells, promote their accumulation, and dampen chronic inflammation.^139^

SMCs were predominant in healthy aortas, and immune cells were predominant in AAAs, which had few contractile SMCs. SMCs transitioned to a synthetic phenotype marked by loss of contractile proteins, increased protease activity, and pro-inflammatory signaling.^43,135^ Phenotypes of synthetic/modulated SMCs were related to matrix repair and calcification in healthy aorta, and to hypoxia adaptation, angiogenesis, and altered vascular tone in AAAs. Adventitial ECs, including stress-adapted and lymphatic/venous subsets that colocalized with CXCL12⁺ fibroblasts, were more abundant in AAAs, likely supporting an increased demand for lymphatic drainage and immune modulation. Fibroblasts were also more abundant in AAAs. AAAs contained two clusters of medial activated/myofibroblast-like cells with wound-healing and fibrotic phenotypes—also present in healthy aorta, likely fulfilling reparative functions—and two clusters of adventitial PI16^+^/CXCL12⁺ universal fibroblasts, likely serving as active organizers of immune–stromal hubs.

The diverse immune infiltrates in AAAs underscore the widespread activation of innate and adaptive immunity. Among macrophages, we identified a resident anti-inflammatory subset, a lipid-associated TREM2⁺/foamy population consistent with the pro-angiogenic, lipid-laden macrophages reported in atherosclerosis and AAA,^91,102,114^ and inflammatory macrophages with increased matrix-degrading activity.^115–117^ Most striking, however, was the accumulation of lymphocytes, rising from 1.4% in controls to nearly half of all AAA cells. B cells encompassed follicular/germinal-center–like, activated, plasma, and proliferating states and colocalized with CXCL12^+^ fibroblasts, activated CD4 and cytotoxic CD8 T cells to form ATLO-like niches. These sites of adaptive immune activation^50,51^ ^111–113,125–127^ show how stromal and immune compartments coevolve in AAA. ATLOs were primarily found in the adventitia, and there was evidence of invasion into the media, as in advanced aneurysmal disease.^136^ The loss of contractile and reparative medial SMCs and phenotypic switching to pathogenic fibroblasts and pro-angiogenic endothelium in AAA likely weaken the aortic wall, resulting in medial degeneration and remodeling of adventitia into an immune-rich niche that fosters ATLO formation.

This study has limitations. First, cell segmentation in AAA is challenging, given the heterogeneity of the aortic wall. Moreover, segmentation with Xenium relies on nuclear staining, which may underestimate large cells, and not all nuclei may be present in a section. Second, Xenium’s gene panel mitigates bias in cell recovery but does not capture all relevant markers, and MAGMA analyses are limited to included loci. Third, our AAA cohort had advanced disease, limiting insight into earlier stages and the temporal dynamics of stromal–immune remodeling. Finally, our study was observational and the findings are correlative. Demonstrating that the *CXCL12*–*CXCR4* axis contributes to AAA will require mechanistic studies, such as a fibroblast-specific *Cxcl12* depletion in inducible mouse models of AAA.

By spatially linking stromal diversity with immune architecture at subcellular resolution, we found that AAA is a disease of coordinated stromal–immune reprogramming and identified the *CXCL12–CXCR4* axis as a potential therapeutic target to mitigate aneurysm progression and rupture risk.

## ACKNOWLEDGMENTS

The authors thank the research participants and organ donor families for their participation.

